# Structural innovation in proteins is constrained by past selection for mutational robustness

**DOI:** 10.1101/2025.08.27.672565

**Authors:** Samuel H. A. von der Dunk, Kamal Dingle, Ard A. Louis, Berend Snel, Paulien Hogeweg

## Abstract

Darwinian evolution has produced two types of functional polymers with different chemical alphabets, RNA and proteins. Dominance of proteins across many cellular functions in extant organisms has been classically attributed to catalytic superiority over RNA. From this perspective, functional RNAs are fossils from the origin of life, where they possibly kick-started molecular evolution. Yet, in addition to direct catalytic capability, the chemical alphabets of RNA and proteins give rise to distinct evolving codes. A crucial question is therefore how the evolution of functional structures is shaped by mutational variation for each of these codes. Here, we employ RNAfold and ESMFold structure predictions to directly compare mutational profiles—*i.e*. the distribution of secondary structures upon point mutations—of functional RNA and proteins. Unlike RNA, natural protein structures feature a narrow mutational profile with high mutational robustness but limited structural variation. Moreover, while functional and random RNA sequences share similar mutational profiles, secondary structures of natural proteins are substantially more robust than those of random sequences and *de novo* proteins, revealing an evolutionary drive toward robustness. Integrating structure prediction with phylogenetic data and simulations, we find evidence for the hypothesis that robust protein structures are favored by selection to buffer errors in translation. As a side-effect, the propensity for structural innovation is reduced in proteins. Our results show that protein folds have evolutionary properties distinct from RNA structures, with implications for the origin of function in both types of molecules—in present-day organisms, as well as during the origin of life.

**Significance statement:** How changes in RNA and protein sequences propagate to their structure through the folding process is fundamental to understanding the evolutionary behavior and functional potential of these polymers. Using computational structure predictions, we argue that protein structures have been selected to exhibit high robustness to translation errors, which, as a side-effect, limits access to structural variants for evolutionary innovation. In stark contrast, RNA structures are innately robust and evolvable, and show no signature of selection. Our work thus shows how mutational effects can be molded by evolution for fixed genotype-phenotype maps. In maps that themselves evolve, such as gene regulatory networks, the imprint of past selection is likely even stronger.

## Introduction

RNA and proteins are the two main catalytic polymers produced by nature. In each case, an analogous folding process turns a specific sequence of monomers into a three-dimensional fold that can supply a catalytic function. Yet, RNA and proteins build upon distinct biochemical rules (Thirumalai and Hyeon, 2005; Rother et al., 2011). RNA folds are largely organized by simple base-pairing of the four nucleotides. In contrast, protein folding is more complex, with a chemically versatile mix of twenty amino acids organizing structure through a mix of electrostatic forces, hydrophobic interactions and hydrogen bonds.

Proteins predominate across many cellular functions in extant organisms, which is generally attributed to catalytic superiority over RNA (Narlikar and Herschlag, 1997; Thirumalai and Hyeon, 2005; Doudna and Lorsch, 2005). Only a few examples of catalytic RNA have been found (Kruger et al., 1982; Guerrier-Takada et al., 1983), most notably ribosomal RNA (rRNA), which catalyses the translation of messenger RNA into protein sequence (Nissen et al., 2000). This central role of RNA in protein production provides striking support for the RNA world hypothesis, which postulates that RNA catalysts preceded proteins during the origin of life (Gilbert, 1986). When proteins arose, they outperformed and replaced most catalytic RNA except rRNA and a few other ribozymes (Jeffares et al., 1998), the reasons for which are still debated (Noller, 2012; Bowman et al., 2020).

Irrespective of catalytic potential, RNA and proteins both face a common problem: the vast majority of sequences do not fold into functional structures (Maynard Smith, 1970; Ellington and Szostak, 1990; Keefe and Szostak, 2001). How evolution traverses sequence space depends on the mapping from sequences, which are subject to mutation, to functions, which are subject to selection. This genotype–phenotype map, as it is more generally known, shapes the course of evolution (Alberch, 1991; Manrubia et al., 2021).

Folding is the physical process that links sequences to structures, and already exposes the impact of a complex genotype–phenotype map on evolution. RNA folding in particular, mapping each RNA sequence to its minimum-free-energy (MFE) secondary structure, has served as a paradigm system in the field (Schuster et al., 1994). Facilitated by efficient structure prediction algorithms (Lorenz et al., 2011), theoretical studies have revealed that RNA structures are both robust and evolvable (Huynen et al., 1996; Huynen, 1996; Wagner, 2005, 2008), consistent with experimental observations (Schultes and Bartel, 2000; Hayden et al., 2011; Jiménez et al., 2013). Sequences encoding the same secondary structure tend to form large neutral networks in sequence space that can be traversed by mutations, granting RNA both immediate robustness and, through neutral sequence divergence, long-term evolvability (Huynen et al., 1996; Huynen, 1996; Wagner, 2005, 2008; Draghi et al., 2010). Mutational robustness and evolvability facilitate structural adaptation and can be further increased by adaptive forces (Wilke, 2001; Sanjuán et al., 2007; Wagner, 2023), phenotypic bias in the sequence–structure map (Dingle et al., 2015; Johnston et al., 2022) and neutral evolution (Van Nimwegen et al., 1999). Similar properties have since been described in other genotype–phenotype maps, including gene regulatory networks (Ciliberti et al., 2007), metabolic networks (Matias Rodrigues and Wagner, 2009; Barve and Wagner, 2013), and simplified models of protein folding (*e.g*. Lipman and Wilbur, 1991; Li et al., 1996; Greenbury et al., 2016), suggesting that robustness and evolvability are fundamental to genotype–phenotype maps in general (Wagner, 2008; Ahnert, 2017).

For proteins, large-scale assessment of fold-level changes across sequence space has not been accessible, with our understanding of protein evolution based instead on biophysical models of folding stability (Bloom et al., 2006; Echave and Wilke, 2017) and experimental fitness landscapes (e.g. Sarkisyan et al., 2016). Genomic data point to widespread structural conservation: many highly divergent sequences retain functional and structural homology (Chothia and Lesk, 1986; Orengo et al., 1994; Illergård et al., 2009; Lesk, 2010; Gabaldón and Koonin, 2013), and in rare cases similar structures arise from apparently independent evolutionary origins (Orengo et al., 1994; Medvedev et al., 2019; Wright, 2025). These observations indicate that, as with RNA, a single protein fold can be encoded by a large number of sequences (Povolotskaya and Kondrashov, 2010). Conversely, examples of proteins adapting towards new structures are relatively rare (*e.g*. Alexander et al., 2009; Dishman et al., 2021), and large-scale clustering of predicted structures has revealed few novel folds (Barrio-Hernandez et al., 2023; Durairaj et al., 2023). *De novo* gene birth from non-coding sequence offers a potential source of structural novelty (Carvunis et al., 2012; Van Oss and Carvunis, 2019). Although the extent of its prevalence remains debated (McLysaght and Hurst, 2016; Weisman et al., 2020), there are some accepted cases based on genomic and functional evidence (Cai et al., 2008; Li et al., 2010; Zhuang et al., 2019). Many protein domains are thought to be ancient (Ranea et al., 2006; Alva et al., 2015), with a substantial number predating the last universal common ancestor (Wehbi et al., 2024; Moody et al., 2024). Whether the lack of observed structural innovation in proteins is a limitation of current comparative genomics methodology, or a result of real evolutionary dynamics is difficult to resolve without probing the protein sequence–structure map directly.

The recent revolution in protein structure prediction (Baek et al., 2021; Jumper et al., 2021; Abramson et al., 2024) now grants that opportunity. We use ESMFold (Lin et al., 2023), a widely adopted algorithm for protein structure prediction directly from single sequences without requiring an alignment (*e.g*. Meier et al., 2021; Xu et al., 2021; Verkuil et al., 2022; Sahakyan et al., 2025), to explore mutational profiles and robustness of protein structures across sequence space, and compare these with the more familiar case of functional RNA. Despite potential limitations of machine learning approaches (Buel and Walters, 2022; Pak et al., 2023; Chakravarty et al., 2024), they capture much of the complexity in the protein structure universe and achieve accuracy far beyond previous methods. We here focus on aggregate statistics across many mutations, and treat predicted structures as a model of the sequence–structure map rather than ground truth, analogous to the use of MFE predictions for RNA.

## Results

### Measuring mutational profiles of RNA and proteins

We analyze the mutational profiles (Fig. 1)—distributions of secondary structures among point mutants— of functional RNA and proteins, focusing on polymers of length *L* = 300 (a typical length for both types; see Fig. S2 for *L* = 100). Naturally occurring sequences were extracted from two large sequence databases, RNAcentral (The RNAcentral Consortium, 2021) and SwissProt (Bairoch and Apweiler, 2000) (see Methods). For each sequence, 100 single nucleotide mutations were sampled, taking transition-transversion bias into account. For proteins, amino acid mutations were sampled using the genetic code, although alternative mutation protocols yielded very similar results (Appendix A). Duplicate mutants and, in the case of proteins, synonymous or truncated mutants were discarded, yielding an average 92 unique mutant sequences per RNA and 65 per protein.

**Figure 1.**
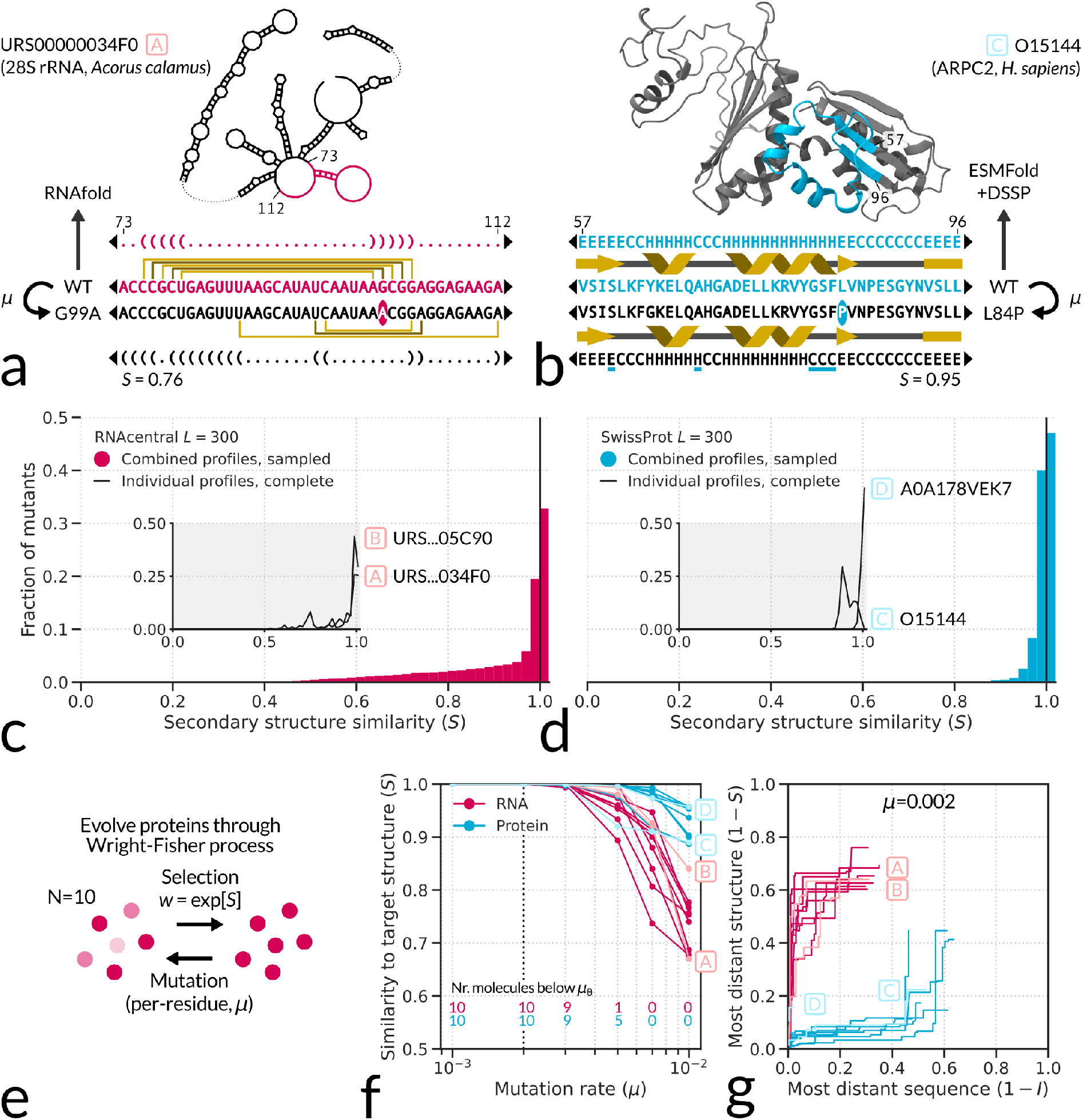
RNA and protein structures exhibit distinct evolutionary signatures and behaviors. (**a**-**b**) We employ a comparative measure (secondary structure similarity, *S*) for the impact of a mutation on molecular structure. Secondary structures of RNA molecules and proteins are represented by linear sequences of three characters, and are predicted by RNAfold for RNA and ESMFold combined with DSSP for proteins. (**c**-**d**) Mutational profiles of natural RNA molecules of length *L* = 300 from the RNAcentral database (*N* = 5451) and natural proteins of length *L* = 300 from the SwissProt database (*N* = 249). The combined profiles are based on up to 100 sampled point mutations per molecule; individual profiles are based on all possible point mutations (900 for RNA, and between 2325 and 2457 for proteins, depending on codon degeneracy). (**e**) Evolutionary simulations with a small number of RNA or protein molecules (*N* = 10), selected for conservation of secondary structure. (**f**) Degree of secondary structure conservation after evolutionary simulation (*i.e*. the highest similarity to the target structure, representing the fittest individual at *t* = 1000), for various mutation rates and across 10 initial RNA molecules and 10 initial proteins (each corresponding to a line). The number of polymers whose target structure is maintained in the simulation and which thus remain below the error threshold is indicated at the bottom of the plot. (**g**) Appearance of novel sequences and secondary structures along the neutral path during evolutionary simulations at low mutation rate, *µ* = 0.002 (big dotted line in panel **f**).

We predicted secondary structures of all natural polymers and their mutant sequences, using RNAfold for RNA (Lorenz et al., 2011), and ESMFold combined with DSSP for proteins (Lin et al., 2023; Kabsch and Sander, 1983, Fig. 1a-b). For both polymer types, secondary structure represents structural conformation along the backbone, coarse-graining the three-dimensional fold to a level where predictions are stable (though our results are not sensitive to this specific representation, see Suppl. Fig. S3, S4). In both cases, secondary structure is also reprented by a string of three characters, facilitating direct comparison (Dingle et al., 2022b). RNA secondary structure is represented with dot-bracket notation—dot for unpaired bases, left and right bracket for paired bases—and protein secondary structure with DSSP notation—`C` for coil, `H` for *α*-helix, and `E` for *β*-sheet. With this classification, our protein structures are made up of 44.0% *α*-helix, 43.7% coil, and 12.3% *β*-sheet.

Following structure prediction, mutant secondary structures are aligned to their respective wildtype and similarity *S* is calculated as the Needleman-Wunsch alignment score divided by polymer length (using 1 for match, 0 for mismatch and −1 for gap; Needleman and Wunsch, 1970). Thus, *S* = 1 corresponds to structural identity, whereas *S* = 0 corresponds to absence of structural similarity. The structural impacts of all sampled mutants are assessed independently and collected in a mutational profile (Fig. 1c-d). We verified that our methodology is reliable by showing that (i) our ESMFold–DSSP combination recapitulates experimental data on structural differences between protein pairs over a large range of sequence differences (Suppl. Fig. S5), (ii) protein structure prediction by OmegaFold yields similar mutational profiles as ESMFold (Suppl. Fig. S6, S7), and (iii) RNA structure prediction by MXfold2, a machine learning algorithm like ESMFold and OmegaFold, also yields similar mutational profiles as RNAfold (Suppl. Fig. S8).

### Secondary structures of proteins are more robust than those of RNA

RNA structures of length *L* = 300 from RNAcentral exhibit a broad mutational profile (Fig. 1c). A sizable fraction of point mutants are neutral (neutrality *λ* = 0.33), providing mutational robustness, in line with previous studies (Schuster et al., 1994; Von der Dunk et al., 2025). At the same time, some mutants fold into very different secondary structures, providing immediate phenotypic variation for adaptation, *i.e*. short-term evolvability.

Protein structures of length *L* = 300 from SwissProt instead exhibit a narrow mutational profile (Fig. 1d), with an even larger fraction of neutral mutants than RNA (*λ* = 0.47, Fig. 1d). Unlike RNA, we find almost no mutants with large structural change, suggesting that protein structures are not as evolvable in the short term. Some individual proteins deviate from the general pattern (e.g. protein O15144 marked as molecule ‘C’ in Fig. 1), showing a mutational profile that is shifted towards lower secondary structure similarities (inset Fig. 1d). Yet even in these cases, mutants with the same extent of structural change as seen in RNA are not observed.

To assess how these mutational profiles impact the evolutionary behavior of our two types of polymers, we performed a set of simple evolutionary simulations (Fig. 1e). In each simulation, a small population of 10 molecules (chosen for computational constraints) is evolved through discrete generations. Each generation, new molecules are selected from the previous population based on their structural similarity to the initial molecule (the “target structure”), calculated from structure predictions in the same way as in our mutational profiling. Sequences of selected molecules are subjected to mutations with per-residue mutation rate *µ* (for proteins, amino acid conversions are weighted by the genetic code), and the cycle is repeated.

Using the above simulation protocol, structural conservation is assayed for a range of mutation rates. At low mutation rates (*µ ≤* 0.002), selection manages to preserve the initial secondary structure in the population for all 20 polymers (Fig. 1f). At high mutation rates, the fittest individual cannot be preserved by selection in our evolutionary protocol, revealing an error threshold (*µ*_θ_, Eigen, 1971) that is surprisingly similar for RNA and proteins. However, while RNA molecules rapidly ‘melt’ to very different structures, proteins remain structurally close to the target, showing the presence of strong structural correlations in protein sequence space.

Below the error threshold (*e.g. µ* = 0.002), populations evolve along the neutral network, exploring sequence space while preserving the target structure (Fig. 1g). Our simulations reveal that RNA and proteins are both able to traverse far into sequence space, but while RNA molecules encounter distinct structures early on (high short-term evolvability), proteins only encounter distinct structures after substantial sequence divergence. These observations again indicate that structural correlations are strongest in protein sequence space. Thus, RNA and protein folding yield distinct evolutionary capacities: proteins exhibit a smoother, less-rugged sequence–secondary structure map than RNA, such that protein structures are more robust but less evolvable than RNA structures.

### Natural protein structures are exceptionally robust to mutations

If the observed mutational profiles of RNA and proteins reflect physical constraints from their folding process and not signatures of evolution, we would expect them to be similar across different phylogenetic domains and to be congruent with random versions of the molecules. To investigate this, we separated our previous natural RNA and protein data by phylogenetic domain, and generated a new random sequence dataset for each polymer type: 1000 random RNA sequences of length *L* = 300 constructed by uniform sampling of nucleotides, and 1000 random protein sequences of length *L* = 300 constructed by uniform sampling of nucleotide triplets after discarding stop-codons.

For RNA, mutational profiles of secondary structures of length *L* = 300 found in bacteria and eukaryotes are extremely similar (Fig. 2a). In fact, even the mutational profiles of secondary structures of random RNA sequences are indistinguishable from those of natural RNA. In line with previous studies (Schuster et al., 1994; Von der Dunk et al., 2025), mutational profiles of natural RNA secondary structures do not carry signatures of evolution. These observations indicate that the evolutionary capacities observed in natural RNA structures, *i.e*. robustness and evolvability, reflect physical constraints from their folding process.

**Figure 2.**
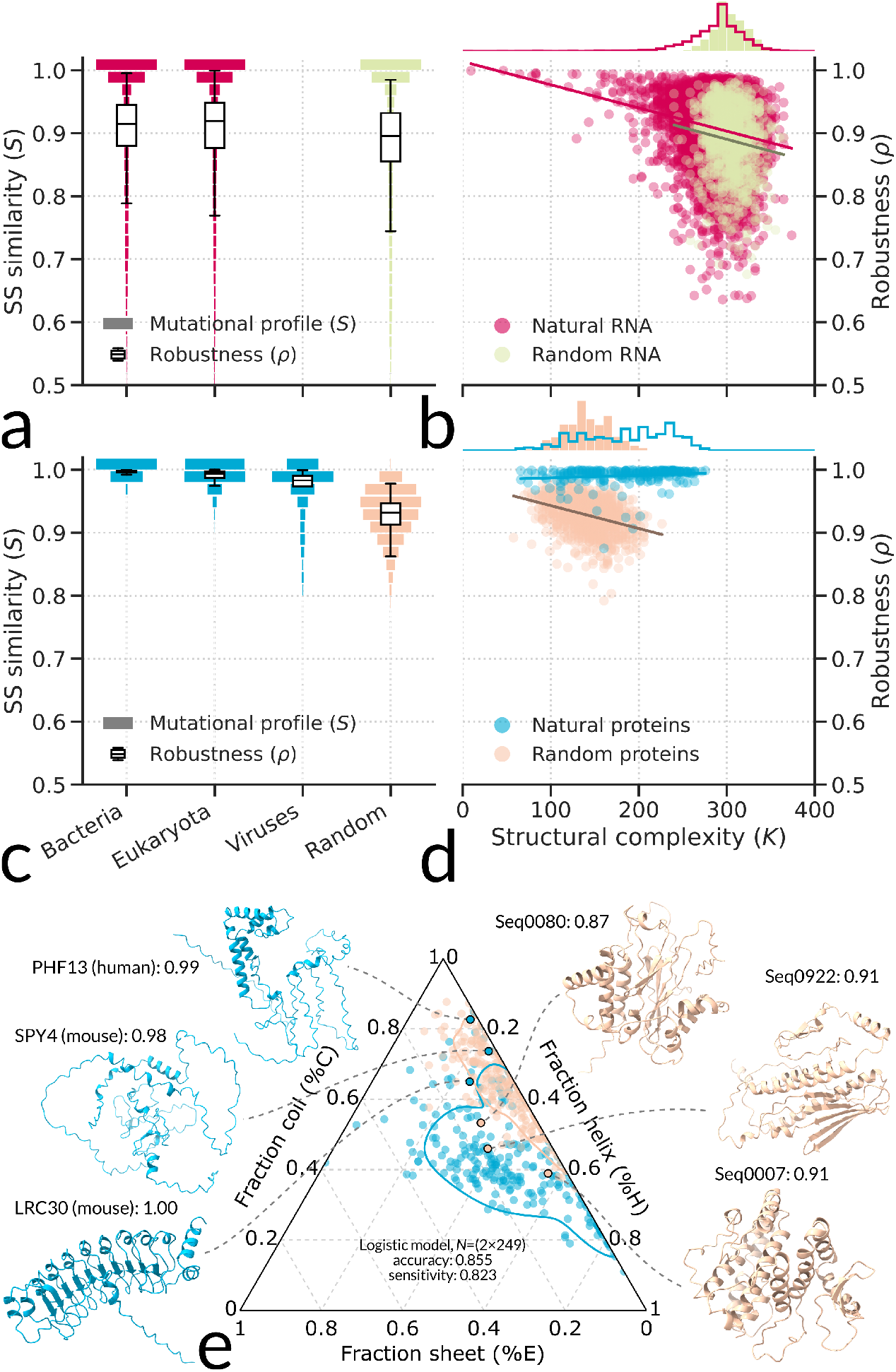
Natural protein structures have evolved exceptional mutational robustness. (**a**) Combined mutational profiles (secondary structure similarity *S*, histograms) and mutational robustness per molecule (mean secondary structure similarity, *ρ*, boxplots) are highly similar across natural and randomly generated RNA sequences. (**b**) Mutational robustness correlates with structural complexity for natural and random RNA (regression slopes *−*3.8 *×* 10^*−*4^ and *−*3.7 *×* 10^*−*4^, both significant with *p <* 0.01). (**c**) Combined mutational profiles (*S*, violins) and mutational robustness per molecule (*ρ*, boxplots) are different for natural and randomly generated proteins. (**d**) Mutational robustness correlates with structural complexity in random but not in natural proteins (regression slopes *−*3.6 *×* 10^*−*4^ and 0.41 *×* 10^*−*4^, significant with *p <* 0.01 and *p* = 0.02). (**e**) Natural proteins show relatively high fractions of *β*-sheet while random proteins show higher fractions of coil. Nevertheless, outliers— *e.g*. natural proteins with high fractions of coil, and random proteins with high fractions of *β*-sheet—still display their characteristic mutational robustness pattern: high robustness for natural proteins and low robustness for random proteins.

For proteins, mutational profiles of secondary structures of length *L* = 300 differ across different phylogenetic domains (*N* = 14, Fig. 2c). Bacterial protein structures (*N* = 76) exhibit higher mutational robustness (measured as mean secondary structure similarity, *ρ*) than eukaryotic protein structures (*N* = 159, Fig. 2c). Viral protein structures, though rare in our data (*N* = 14), showed even lower mutational robustness. Hence, mutational profiles are different in phylogenetic domains with different macro-evolutionary patterns.

Secondary structures of random protein sequences show markedly lower mutational robustness than those of natural proteins. In fact, only a tiny fraction *λ* = 0.004 of sampled mutations of random proteins was neutral at the secondary structure level. Thus, proteins are not inherently robust to mutations and high robustness of natural proteins appears to be an acquired property. Unlike RNA, mutational profiles of natural proteins carry a clear signature of evolution.

Previous work has shown that RNA structures with high frequency in sequence space are more robust (Greenbury et al., 2016; Mohanty et al., 2023) and less complex as measured by compression of their secondary structure representation (Dingle et al., 2018; Von der Dunk et al., 2025, see Methods).

Together, these points suggest that a negative correlation exists between mutational robustness *ρ* and structural complexity *K*, and our analysis indeed uncovers such relationship for RNA (Fig. 2b). We find that random proteins harbor the same negative correlation between mutational robustness and structural complexity (Fig. 2d). However, natural proteins do not: those with high structural complexity still exhibit exceptionally high robustness. Interestingly, we find that some natural RNA structures are much simpler than random RNA structures (especially RNA molecules with repetitive sequences, see Von der Dunk et al., 2025), but that many natural protein structures are substantially more complex than random protein structures (see marginal distributions in Fig. 2d). Taken together with their exceptionally high mutational robustness, natural proteins with complex structures are particularly well separated from the bulk of protein space represented by random sequences.

Next, we investigated differences between natural and random protein structures, asking if these could explain the large difference in mutational robustness between natural and random proteins. To this end, we characterized protein structures through their composite fractions of coil, *α*-helix and *β*-sheet, revealing a clear separation of natural and random protein structures (Fig. 2e). ESMFold predicts natural proteins to contain larger proportions of *β*-sheet and random protein structures to contain larger fractions of coil. We can even use this information to predict the source of a protein structure (natural versus random) with an accuracy of 0.855 and a sensitivity of 0.823 by logistic regression. Still, these structural differences do not explain the disparity in mutational robustness between natural and random proteins. For example, three natural protein outliers that resemble random protein structures (high fraction of coil, low fraction of *β*-sheet) show remarkably high robustness. Conversely, three random protein outliers that resemble natural protein structures (low fraction of coil, high fraction of *β*-sheet) and which even appear to form compact globular folds, show remarkably low robustness. Thus, the data suggest that natural proteins have been under evolutionary pressure to increase mutational robustness independent of their specific structure.

### Natural proteins are structurally diverse

The exceptionally high mutational robustness predicted for natural proteins could suggest that they occupy a limited region of sequence and structure space. On average, two natural proteins share approximately 10% of their sequences, similar to two random proteins (Fig. 3a). Pairs of natural proteins also share approximately 30% of their secondary structures, which, surprisingly, is much less than two random proteins (Fig. 3a). We already observed that natural proteins occupy a much larger range of structural complexity than random sequences (Fig. 2d), which is opposite from natural RNA whose structural complexity distribution is very similar to that of random RNA (Fig. 2b, Von der Dunk et al., 2025). Going further, we observe higher diversity of natural proteins across 7 secondary structure features (*i.e*. the fraction and number of contiguous segments of each type of secondary structure, and structural complexity) organized through principal component analysis (Fig. 3c-d). All these observations suggest that high mutational robustness does not constrain protein structure diversity. Evolution has shaped extant proteins to display a wide range of structural phenotypes compared to random sequences, in addition to their high robustness.

**Figure 3.**
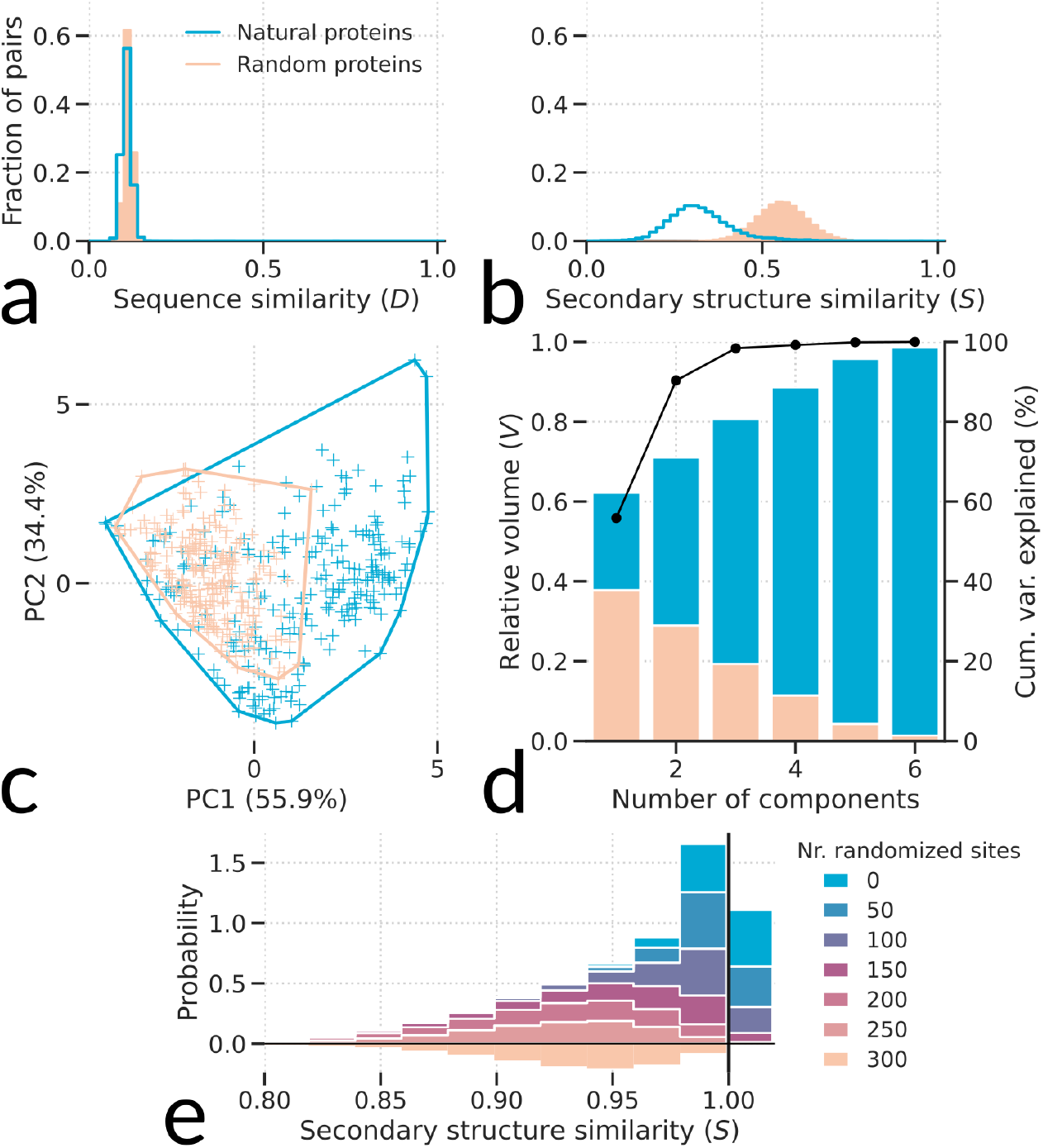
Natural proteins are structurally more diverse than random proteins. (**a**) Sequence similarity and (**b**) secondary structure similarity between all pairs of natural proteins, and between all pairs of random proteins. (**c**-**d**) Natural proteins show larger diversity in every structural dimension (see text), organized by principal component analysis. (**c**) Occupied volumes of natural and random proteins in the first two principal components. (**d**) Relative high-dimensional occupied volumes when adding more principal components. (**e**) Natural proteins are far removed from random sequences: mutational profiles of natural proteins after accumulating mutations, shown per 50 sites.

To find out on what timescale random proteins could potentially evolve high mutational robustness characteristic of natural proteins, we measured the proximity between natural and random proteins in sequence and structure space. We performed several iterations in which natural proteins were mutated at 50 additional sites, *i.e*. resulting in natural proteins with 50, 100, 150, 200 and 250 sites mutated (Fig. 3e). It takes as many as 150 mutations (*i.e*. randomizing half of the protein sequence) to arrive at a mutational profile that resembles that of random proteins—in particular, where neutral mutants are rarely observed anymore. Note that this number is consistent with our previous evolutionary simulations at low mutation rate, where natural proteins encountered distinct structures only when around half of their sequence was mutated (Fig. 1g). While these 150 mutations do not necessarily represent the shortest mutational path, it does suggest that, on average, extensive sequence change is needed to erase the distinctive robustness signature of natural proteins.

### *De novo* yeast proteins resemble random sequences

Our results seem to suggest that substantial evolution is required to increase the mutational robustness of proteins relative to that of randomly generated and non-evolved sequences. Following this reasoning, we might ask how robust *de novo* proteins are which have a hypothesized recent origin from non-coding DNA (Levine et al., 2006; Begun et al., 2007). *De novo* proteins were once thought to be rare, but as more curated candidates are being identified, *de novo* gene birth may actually represent an important route for the origin of new proteins (Magadum et al., 2013; McLysaght and Guerzoni, 2015). Thus, we now turn to the *Saccharomyces cerevisiae* proteome, a large and diverse set of proteins with a length range of 50 *≤ L ≤* 2014 (the upper boundary being the limit for ESMFold) and a small number of annotated *de novo* proteins (Vakirlis et al., 2020). These *de novo* annotations are based on a combination of protein-coding signatures (*i.e*. the lack thereof) and phylogenetic analysis (*i.e*. absence of homologs outside the *Saccharomyces* genus). In the Appendix, we confirm our findings as described below with proteome analyses of *Oryza sativa* and *Drosophila melanogaster* for which *de novo* annotation is also available (Suppl. Fig. S9).

Established (*i.e*. non-*de novo*) yeast proteins, ranging more than an order of magnitude in length, feature similar high mutational robustness as natural proteins of length *L* = 300 (Fig. 4a). In contrast, *de novo* proteins vary much more in robustness, with 51/188 (27%) that display robustness similar to that of established proteins (*ρ* within two standard deviations) and 137/188 (73%) displaying lower robustness, similar to random sequences. Moreover, *de novo* proteins generally follow the robustness versus complexity correlation—complexity now normalized by length—that was also observed for random proteins of length *L* = 300 (Fig. 4b), unlike established proteins. Thus, we find that most proteins that recently emerged from untranslated DNA are not yet robust and still resemble random proteins in that respect, in agreement with experimental findings (Edgell et al., 2020).

**Figure 4.**
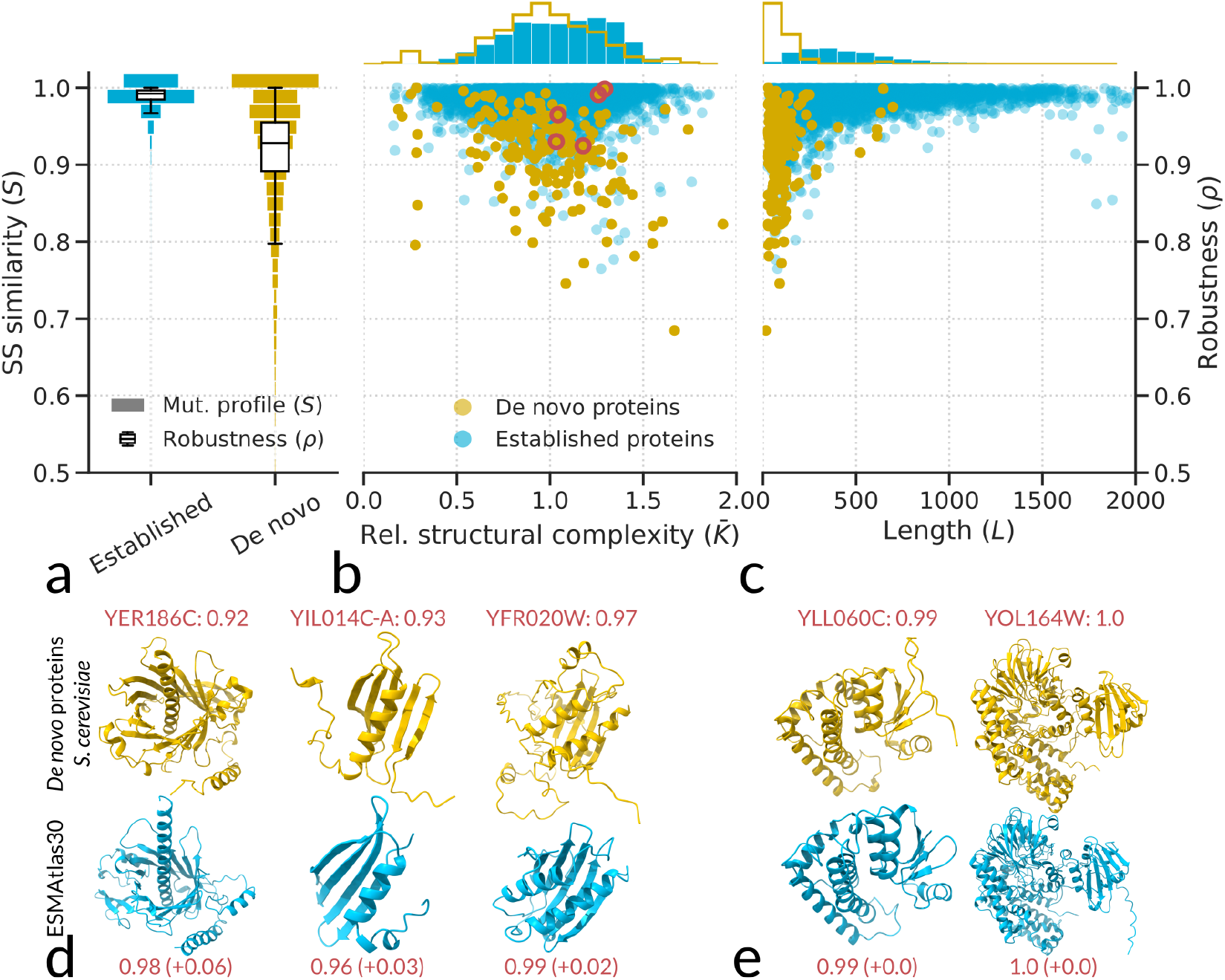
*De novo* proteins in *Saccharomyces cerevisiae* predicted by Vakirlis et al. (2020) exhibit low mutational robustness. (**a**) Combined mutational profiles (*S*, violins) and per-molecule means (*ρ*, boxplots) for *de novo* (*N* = 188) and established (*N* = 5217) proteins from *S. cerevisiae*. (**b**) Mutational robustness decreases with structural complexity in *de novo* but not established proteins (regression slopes *−*0.070 and 0.006, both significant with *p <* 0.01). (**c**) Mutational robustness decreases with molecule length across all *S. cerevisiae* proteins (regression slope 1.67, significant with *p <* 0.01). (**d**) Proteins from the ESMAtlas30 database with structural similarity to *de novo* proteins as detected by FoldSeek have higher robustness (*ρ*, values indicated in red). (**e**) Our analysis highlights two suspicious *de novo* proteins with high mutational robustness and high structural complexity, `YOL164W` and `YLL060C` (see panel **b**), for which sequence homology searches point to horizontal gene transfer instead of *de novo* origin.

To see if *de novo* proteins have distinct structural characteristics that explain their low robustness, we searched for structural analogs in the ESM Metagenomic Atlas using FoldSeek (Lin et al., 2023; Van Kempen et al., 2024). Three of our 188 *de novo* proteins produced a significant hit (E-value *<* 0.05) with high structural similarity (Fig. 4d). In all three cases, the structural analog is more robust to mutations than the *de novo* protein, indicating that there are no inherent physical constraints that prevent these *de novo* structures from achieving higher robustness. Rather, it appears that *de novo* proteins have not had enough time to evolve high mutational robustness to the same level of older established proteins. For two additional *de novo* proteins (`YOL164W` and `YLL060C`), we found that the highly significant bacterial hits were in fact homologs (Fig. 4e; pHMMer log(*E*) *< −* 100, see Suppl. Data S1, S2), indicating that these two proteins likely originate from horizontal gene transfer and are wrongly annotated as *de novo* proteins. This conclusion is supported by their combination of high predicted robustness and high structural complexity, something that is rare for random and *de novo* proteins (Fig. 4b). Thus, while it remains extremely challenging to infer *de novo* proteins, analysis of mutational robustness and complexity could provide independent evidence for such annotations.

*De novo* proteins are known to be relatively short, and we find that their length partly explains their low robustness (Fig. 4c, Suppl. Fig. S9a). On average, short proteins (*L <* 250) show lower mutational robustness even in putative established proteins. Short proteins also feature a broader mutational profile because the relative impact of a point mutation is larger, and there is an increased probability that the structural conformation of all residues is conserved (Suppl. Fig. S2). Short proteins are more evolvable, resembling RNA, and could present a suitable starting point for the evolution of novel structures. Conversely, protein length and mutational robustness increase with age (Suppl. Fig. S10).

### Mutational robustness buffers translational errors

We next investigated the potential evolutionary drivers of mutational robustness in natural protein structures. Based on the sequence–structure–function paradigm, we would predict that proteins with higher mutational robustness can accommodate more sequence divergence over evolutionary time while still conserving structure and function. Sequence divergence has been studied extensively in yeast (Pál et al., 2001; Wall et al., 2005; Drummond et al., 2005a, 2006; Serohijos et al., 2012, 2013), allowing us to explore the correlation with our mutational robustness predictions (Fig. 5).

**Figure 5.**
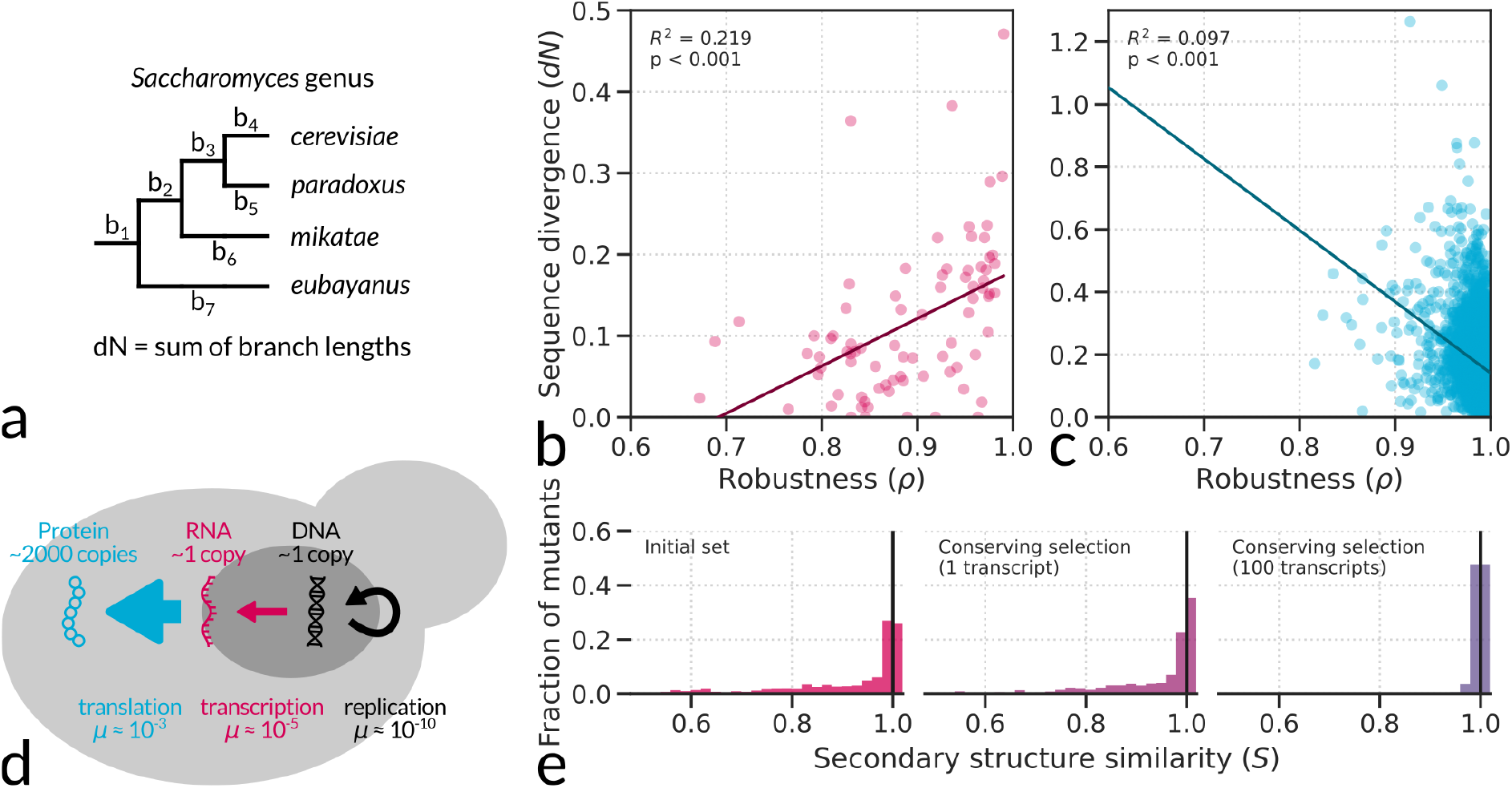
Negative correlation between sequence divergence and robustness in the *S. cerevisiae* proteome, points to translational errors as a driver in the evolution of mutational robustness. (**a**) Sequence divergence was inferred using CodeML (Yang, 2007). (**b**-**c**) Opposite correlations between sequence divergence and mutational robustness in RNA molecules (*N* = 81, regression slope 0.583) and proteins (*N* = 4082, regression slope *−*0.311) from *S. cerevisiae*. (**d**) Schematic showing typical copy numbers of molecules inside *S. cerevisiae* and error rates in the corresponding production processes (Milo and Phillips, 2015). (**e**) Mutational profiles of 10 natural RNA molecules of length *L* = 300 (same as in Fig. 1) before (left panel), and after evolutionary simulation (*t* = 10^4^), with conserving selection acting on the secondary structure of a single transcript (middle) or on that of 100 transcripts for each molecule (right).

Following previous studies, we calculated sequence divergence for a large number of established proteins as the rate of non-synonymous substitutions *dN* between four closely related yeast species—*S. cerevisiae, S. paradoxus, S. mikatae* and *S. eubayanus* (Fig. 5a). Drummond et al. (2006) pinpointed gene expression level as the single strongest correlate with sequence divergence in proteins, accounting for 28.8% of variation. We here find that mutational robustness is able to explain a small but significant fraction of 9.7% of variation in sequence divergence of proteins (Fig. 5c). In contrast to RNA though (Fig. 5b), the correlation between mutational robustness and sequence divergence is negative rather than positive: proteins with the most robust structures show the least sequence divergence. The negative correlation between mutational robustness and sequence divergence suggests that highly expressed proteins which, as pointed out by Drummond et al. (2006) evolve slower at the sequence level, also experience stronger selection for mutational robustness of their structure. A possible explanation for this is that errors during translation of highly expressed proteins pose a burden for the cellular machinery that protects against misfolded proteins, and thereby could drive selection for mutational robustness. Given that errors in translation occur at much higher rates than errors in replication (Mordret et al., 2019), the former may in fact be what drives evolution towards stable protein structures with limited mutational access to structural variation (Fig. 5d).

To test the hypothesis that proteins face *in vivo* selection against errors through their high copy numbers, we performed two *in silico* evolution experiments with RNA molecules (Fig. 5e). As before (cf. Fig. 1e), we evolve populations of RNA molecules (though larger populations, *N* = 100) for 1000 generations starting with one of the natural *L* = 300 RNAcentral molecules and selecting for the initial structure as the target. In the first experiment, selection is on each molecule directly; in the second experiment, 100 copies are created from each molecule applying a per-nucleotide error rate equal to the mutation rate, and selection is on the average similarity of those 100 copies to the target structure. We run 3 replicates for each of 10 different natural RNA molecules in each experiment.

Initially, the 10 natural RNA structures show broad mutational profiles as described before (cf. Fig. 1c), except with more noise due to the small number of assessed molecules. When molecules evolved directly under conserving selection, the mutational robustness and fraction of neutral mutants are slightly increased, in line with previous theoretical work (Van Nimwegen et al., 1999). When molecules evolved under conserving selection while undergoing translational amplification, the mutational robustness and fraction of neutral mutants are increased to an even higher degree, resembling mutational profiles of natural protein structures (Fig. 5d). Thus, simple evolutionary simulations align with phylogenomic data from *S. cerevisiae* and suggest that translational errors pose a crucial selective constraint on proteins but not on RNA, explaining differences in their evolved evolutionary properties reflected in their mutational profiles.

## Discussion

Our main finding is that RNA molecules and proteins exhibit distinct evolutionary properties through their sequence–secondary structure map. For RNA, folding yields structures with innate robustness and evolvability. For proteins, folding by default yields structures with low robustness, but this can be overcome by evolutionary processes to build highly robust folds. We further suggest that such highly robust protein folds result from selection for buffering of translational errors. Functional RNAs, by contrast, do not experience the same translational amplification, and being robust by default, exhibit no detectable evolutionary signatures.

Our framework employs different types of structure prediction algorithms for RNA and proteins. Despite the fact that ESMFold and related machine-learning-based algorithms have revolutionized protein structure prediction, it is difficult to know how well they extrapolate to sequences that do not resemble their training data, such as random sequences. Instead, RNAfold is based on thermodynamics and likely generalizes well. However, we obtain the same broad mutational profiles for RNA secondary structures when we use the machine-learning-based algorithm MXfold2 to predict those structures, indicating that these algorithms can in principle ‘learn’ molecular folding in sufficient detail to infer aggregate statistics such as robustness and evolvability.

Extraction of protein secondary structure from tertiary structure provides another way to reduce uncertainty of prediction algorithms. At the level of secondary structure, different algorithms consistently predict narrow mutational profiles with high robustness for natural proteins (Suppl. Fig. S6, see also Dingle et al., 2022b). For random sequences, mutational profiles still vary between algorithms. While state-of-the-art tertiary structure prediction algorithms ESMFold, OmegaFold and AlphaFold consistently predict low mutational robustness, secondary structure prediction algorithms predict higher mutational robustness, closer to natural proteins. We think that the low sensitivity to mutational effects of these secondary structure predictors may be due to the low level of detail in their trained sequence–structure mapping. Even if coarse-graining to secondary structure provides a more informative (less noisy) prediction in the end, tertiary structure is vital for learning the physics of protein folding.

RNA has been dubbed an “ideal evolvable molecule”, combining mutational robustness with evolvability of structure (Schuster et al., 1994; Hogeweg, 2012). The striking similarity between natural and random RNA structures has been highlighted before (Fontana et al., 1993; Smit et al., 2006; Jörg et al., 2008; Stich et al., 2008; Dingle et al., 2015, 2022a; Von der Dunk et al., 2025). It suggests that function may readily appear in a pool of random RNA molecules, for instance during the origin of life. Recent discovery of a small auto-catalytic RNA molecule lends further support to the RNA world hypothesis (Gianni et al., 2026).

Our findings show that proteins contrast sharply with RNA, with clear differences between natural proteins and random sequences, aligning with two previous studies (*e.g*. De Lucrezia et al., 2012; Ferrada and Wagner, 2012), although another study instead highlighted similarities (Tretyachenko et al., 2017). Protein structures face a tradeoff between mutational robustness and mutational variation, captured in their narrow mutational profile. As a consequence of this tradeoff, the two main routes to new proteins, gene duplication and *de novo* gene birth, both face unique challenges. Gene duplication creates a copy of a pre-existing robust structure with limited access to new structures through mutation. As our simulations have shown, substantial sequence divergence is required to evolve a new structure. As a result, folds of duplicated proteins may be more likely to be conserved and repurposed for a new cellular niche rather than being substantially modified (Cardarelli et al., 2010). In contrast, *de novo* gene birth directly creates novel structures with more access to new structural variants through mutation than established proteins. However, these *de novo* structures feature low mutational robustness and are therefore hard to preserve over evolutionary timespans.

One way to avoid the tradeoff between robustness and evolvability in protein structures is to start with a short protein, as in the case of *de novo* and unconserved proteins in *S. cerevisiae* (Lipman et al., 2002). Short proteins have a relatively broad mutational profile compared to longer proteins (see Suppl. Fig. S2), which allows them to navigate the sequence–structure map more efficiently, similar to RNA secondary structures. Once a suitable structure is found, protein length may be increased to enhance mutational robustness. Interestingly, this scenario fits with a recent study on evolutionary simulations of short random sequences selected for stability (Sahakyan et al., 2025), where stable proteins of length *L ≈* 50 rapidly evolved from short peptides of length *L* = 12. Large mutations such as partial duplications played an important role, allowing evolution to duplicate stable substructures to form complete globular domains. In the field of protein design, large mutations are also leveraged to efficiently search the enormous sequence space without getting stuck in local optima (Drummond et al., 2005b; Romero and Arnold, 2009; Fram et al., 2024).

Despite potential alternative routes towards functional proteins, the evolutionary constraints faced by proteins make them an unlikely candidate for the earliest functional biopolymer. The more accessible sequence–structure map of RNA supports an RNA-first scenario in which proteins only emerged when catalytic sophistication became instrumental.

Finally, our findings indicate that intuitions drawn from the RNA sequence-to-structure map do not readily apply to protein secondary structures, in contrast to claims that these principles hold for a wider set of genotype–phenotype maps (Hogeweg, 2012; Wagner, 2013; Manrubia et al., 2021). RNA folding and protein folding both constitute genotype–phenotype maps whose organization is fixed by physics. While innate biases remain reflected in natural RNA structures, we found that selection alters the local topology of the genotype–phenotype map for protein structures. Such an effect or side-effect of past evolution on future trajectories, also dubbed evolution of evolvability (Wagner and Altenberg, 1996; Pigliucci, 2008), likely plays an even bigger role in other genotype–phenotype maps which are not ‘physical’. In metabolic networks (Barve and Wagner, 2013), gene regulatory networks (Crombach and Hogeweg, 2008), and developmental programmes such as morphogenesis (Salazar-Ciudad and Jernvall, 2010), the genotype–phenotype map is itself constructed by evolution, and mutational profiles may promote or hinder phenotypic innovation in surprising ways. In sum, evolutionary dynamics depend on the details of the specific genotype–phenotype map, but also help shape those details, revealing that contingency plays an important role in the evolutionary path from the origin of life towards present-day complexity.

## Methods

### Data preparation

High-quality, natural RNA molecules of length *L* = 300 were selected from RNAcentral (The RNAcen-tral Consortium, 2021). RNAcentral assigns quality control warnings to sequences that are potentially incomplete or not non-coding. Subsequently, sequences were passed through a similarity filter, where every next sequence was only kept in the dataset if it did not share more than 80% identity with any of the already selected sequences. This step was taken to ensure that the natural RNA dataset was not dominated by many copies of very similar RNA sequences. Finally, in line with previous work (Von der Dunk et al., 2025), the few archaeal RNA sequences were discarded, yielding a dataset with *N* = 5451 bacterial and eukaryotic RNA sequences.

High-quality, natural protein sequences were downloaded from UniProt, selecting all reviewed (Swis-sProt) entries of length *L* = 300 (*N* = 981, Ahmad et al., 2025). Similar to RNA, these protein sequences were passed through a similarity filter, keeping only protein sequences with less than 80% similarity to any other protein in the dataset (*N* = 249).

Random RNA and protein sequences were constructed as described in the Results. For proteins, we also tested alternative random sequence sets, *i.e*. sequences created from uniform sampling of amino acids and sequences created by shuffling natural protein sequences, but these even showed lower robustness than protein sequences created from random nucleotide codons (Appendix A).

*S. cerevisiae* protein-coding genes were downloaded from the Saccharomyces Genome Database (SGD Cherry et al., 2012), as nucleotide sequences. Proteins not annotated by Vakirlis et al. (2020) were removed from the dataset. Mutations were performed directly on these nucleotide sequences, discarding sequences with a stop-codon.

For evolutionary sequence divergence (Fig. 5), RNA genes were identified in the four yeast genomes using Infernal (Nawrocki and Eddy, 2013) run with all available RFAM profiles (Ontiveros-Palacios et al., 2025). Protein-coding genes were directly downloaded from SGD, as for *S. cerevisiae*. For identification of orthologous groups, OrthoFinder (Emms and Kelly, 2019) was employed for both RNA and proteins, choosing multiple sequence alignment for gene tree inference, DNA input for RNA genes, and default options otherwise.

### RNA and protein folding

RNA sequences were folded using RNAfold (Lorenz et al., 2011), which predicts the minimum-free energy (MFE) secondary structure, *i.e*. the most probable structural state of the sequence. Secondary structures are uniquely represented in dot-bracket notation (see Fig. 1a). This approach maps, at any given length, 4^*L*^ unique sequences to their corresponding secondary structure. Dot-bracket notation uses three characters, but physical constraints in base pairing bring down the total number of secondary structures to ~ 1.8^*L*^, rather than 3^*L*^ (Dingle et al., 2015). These numbers reveal an enormous redundancy in the genotype–phenotype map, whereby many sequences map to the same secondary structure. The numbers do however not reveal where in sequence space those neutral sequences are found, such as the neutral networks.

Protein sequences were folded using ESMFold (Lin et al., 2023), which predicts three-dimensional structure from which secondary structure elements can be extracted using DSSP (Kabsch and Sander, 1983). Protein secondary structure is represented by DSSP notation (see Fig. 1b). DSSP notation also uses three characters, but these occur at different frequencies (effective number of characters 2^*H*^ = 2.67 for natural proteins, compared to 2^*H*^ = 2.97 for natural RNA), reflecting physical constraints. Interestingly, sequence space of proteins is substantially larger than RNA (20^*L*^ versus 4^*L*^), indicating that the redundancy for protein structures is also much larger (going from 20^*L*^ to *<* 3^*L*^).

### Calculating structural complexity

To calculate structural complexity of an RNA molecule, the dot-bracket notation for secondary structure is converted into a binary string: ‘(‘*→* ‘10’, ‘)’ *→* ‘01’, ‘.’ *→* ‘00’. The binary string is then compressed using Lempel-Ziv compression (Lempel and Ziv (1976) as detailed in Dingle et al. (2018)), with the resulting number of bits representing the complexity of the original object. This measure of complexity originates from the field of algorithmic information theory, and has found a valuable application in RNA folding due to the existence of a phenotypic bias that allows an upper limit of the neutral set size of RNA phenotypes to be predicted from their structural complexity (Johnston et al., 2022).

Following Dingle et al. (2022b), structural complexity of proteins is calculated in a similar way to RNA, by converting the DSSP notation into a binary string: ‘H’ *→* ‘01, ‘E’ *→* ‘10’, ‘C’ *→* ‘00’. The resulting binary string is compressed using the Lempel-Ziv algorithm (as discussed above), with the resulting number of bits representing the complexity of the protein secondary structure. Structural complexity provides a coarse-grained measure of protein structure and a single dimension along which sets of protein structures can be easily compared.

## Data availability

Data and codes are publicly available at Dryad, DOI: `10.5061/dryad.1jwstqk99`.

## Code availability

Data and codes are publicly available at Dryad, DOI: `10.5061/dryad.1jwstqk99`.

