## Supplementary Materials for "Structural innovation in proteins is constrained by past selection for mutational robustness"

November 13, 2026

### Appendix A: The genetic code promotes robustness only slightly

The genetic code impacts the relative abundance of amino acids—by the number of codons assigned to each—as well as their mutational linkage—by the similarity between codons of different amino acids. Previous work suggests that the standard genetic code promotes mutational robustness of protein structures by limiting transitions between residues with very different polar requirement (Freeland and Hurst, 1998). Our framework grants an opportunity to test this hypothesis by using protein structure prediction to measure the impact of the genetic code on mutational robustness (Fig. S1).

First, we created two two additional sets of "random sequences" of length $L = 300$: (i) sequences with uniform random sampling of amino acids, which lack the redundancies of the genetic code, and (ii) shuffled natural protein sequences–two scrambled versions for each natural protein, which preserve the redundancies of the genetic code and additionally capture potential signatures of evolution. Second, we used two additional mutation schemes to sample mutations and measure robustness: (i) uniform random amino acid substitutions, destroying the mutational linkage of the genetic code, and (ii) amino acid transitions as observed in closely related proteins defined by the PAM1 substitution matrix (Dayhoff, 1972), which capture the linkage of the genetic code as well as the effects of other mutational biases and selection.

To facilitate comparison between 12 mutational profiles, we focus on their means and standard deviations (Fig. S1h). These two measures are related across sequence sets. In natural proteins, the mutational profiles are squeezed against the upper limit in secondary structure similarity (*i.e.* S = 1), resulting in high $\rho$ and corresponding low $\sigma$.

The structures of proteins constructed from random codons, as used in the main text, display slightly higher mutational robustness than those constructed from uniform random amino acid sequences, indicating that the genetic code promotes robustness by directing evolution towards a particular set of sequences (Fig. S1a–d,h). However, as the structures of proteins constructed by shuffling natural protein sequences are on average slightly less robust than those constructed from random nucleotide sequences, it is not just the amino acid frequency distribution of natural proteins that explains their high mutational robustness. For instance, a high frequency of aspartic acid (D) and glutamic acid (E) relative to random nucleotide sequences (compare red bars in Fig. S1b–c) is not guaranteed to increase robustness, even though it apparently does so in the specific context found in natural proteins.

In line with previous work (Freeland and Hurst, 1998), the genetic code buffers against structurally harmful mutations (Fig. S1e–h). Non-synonymous nucleotide mutants, as used in the main text, on average share more similar secondary structures with the wildtype than random amino acid mutants do. Nevertheless, the effect of the genetic code in linking structurally similar amino acids is very small (*e.g.* increasing $\rho$ from 0.92 to 0.93 in random codons) compared to the effect of the genetic code in sampling more robust genotypes in the first place (*e.g.* increasing $\rho$ from 0.90 to 0.93 from uniform random amino acid sequences to random codon sequences). The application of PAM1 transition probabilities increases robustness slightly further, suggesting that some of the possible non-synonymous sequence variations have been selected out from the PAM1 homologs based on their deleterious effect on structure. In sum, while there are small effects of the genetic code on mutational robustness of protein structures, these are far from sufficient to explain the high mutational robustness demonstrated by natural proteins.

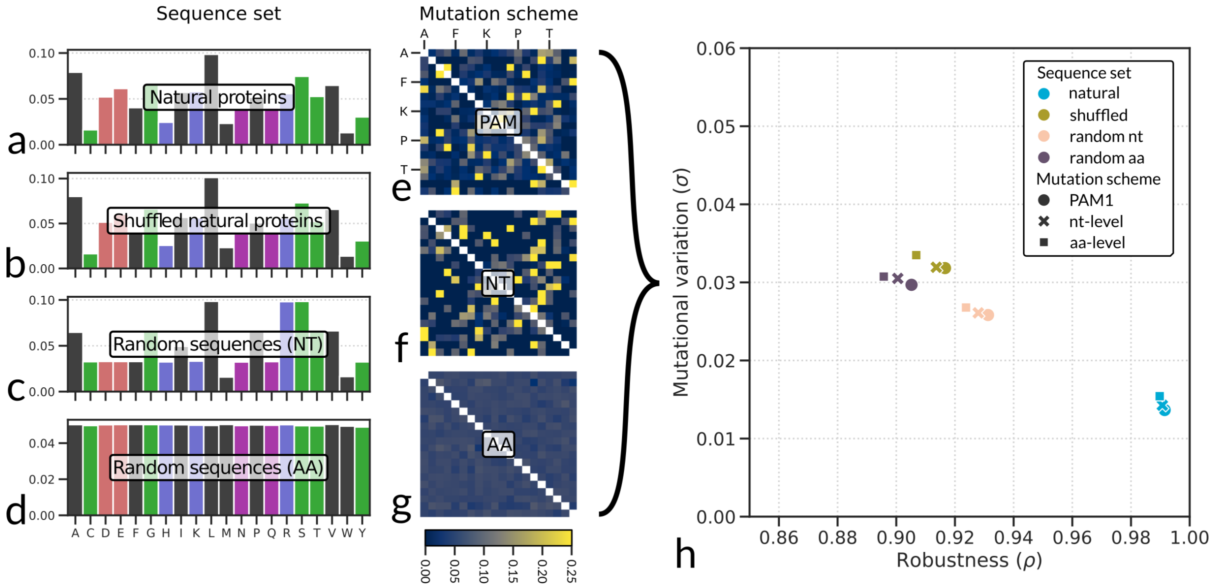


Figure S1 – The genetic code favors mutational robustness in protein sequences of length $L = 300$. (**a**–**d**) Frequency distributions of amino acids in natural proteins and different random sequence sets. (**e**–**g**) Amino acid substitution rates realized by different mutational schemes. (**h**) Comparison of mean ($\rho$) and standard deviation ($\sigma$) of mutational profiles from all 12 mutation assays (4 sequence sets times 3 mutation schemes).

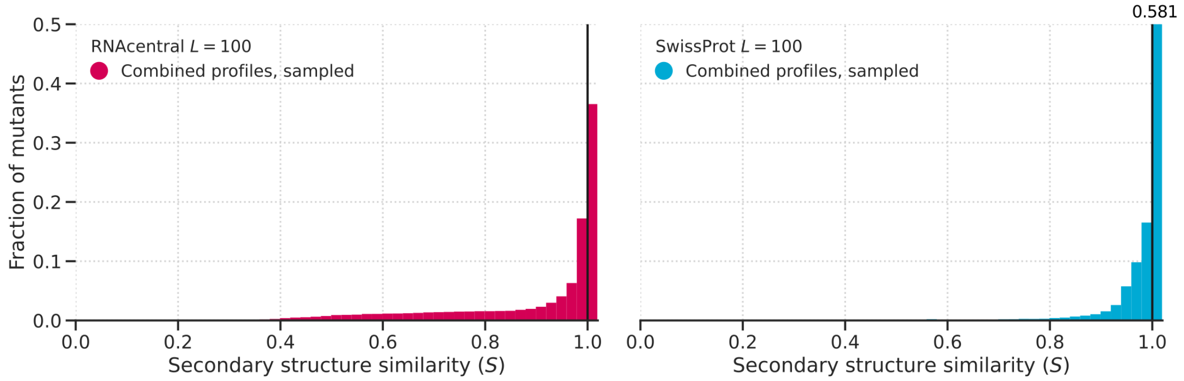


Figure S2 – Mutational profiles of natural RNA molecules of length $L = 100$ from the RNAcentral database ($N = 6094$) and natural proteins of length $L = 100$ from the SwissProt database ($N = 255$). The combined profiles are based on up to 100 sampled point mutations per molecule. Note that both profiles are slightly broader than those inferred at $L = 300$ (Fig. 1c,d in the main text), with a larger fraction of neutral mutants as well as mutants with more severe structural change.

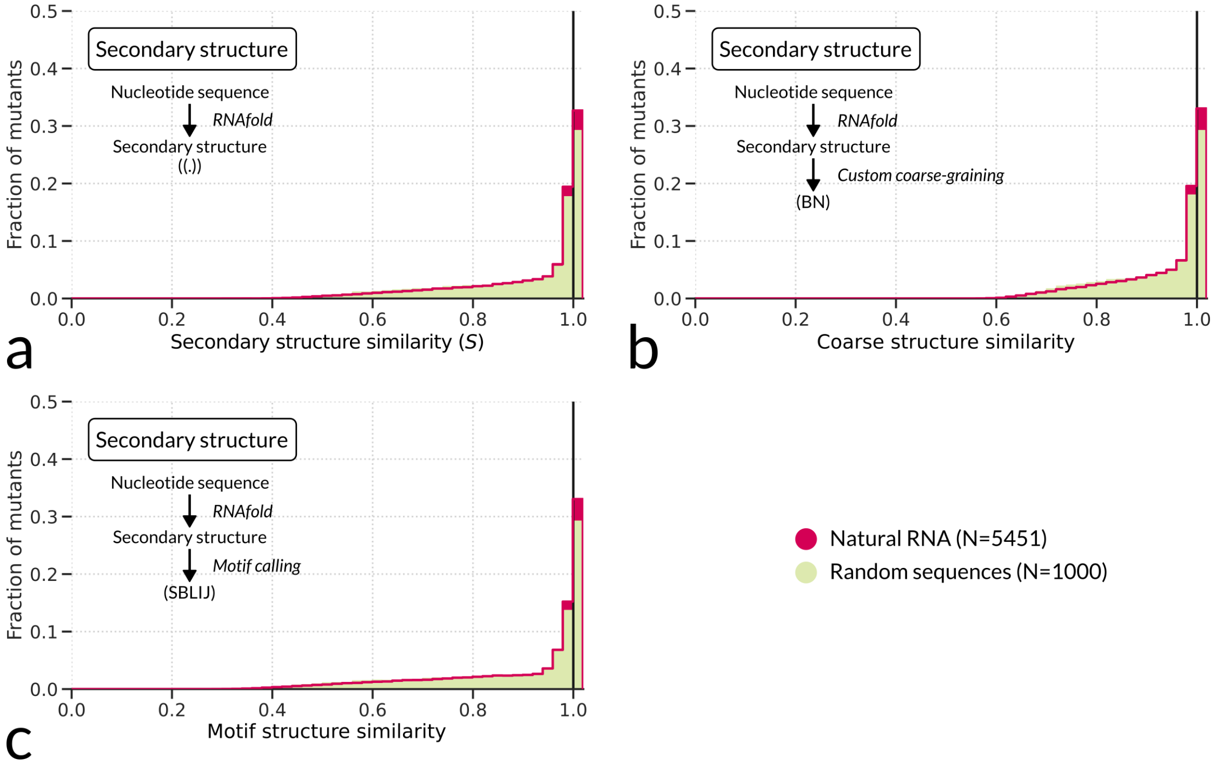


Figure S3 – Mutational profiles of natural and random RNA molecules of length $L = 300$ using different structural representations: (**a**) standard dot-bracket notation, (**b**) coarse-grained representation wherein brackets are combined into character '`B`' and dots are represented with '`N`', and (**c**) motif representation, wherein residues are annotated by their structural motif ('`S`': stack, '`B`': bulge, '`L`': loop, '`I`': internal loop, and '`J`': junction). More detailed structural representations yield slightly larger mutational effects, but the characteristic broad mutational profile of RNA is maintained at all levels.

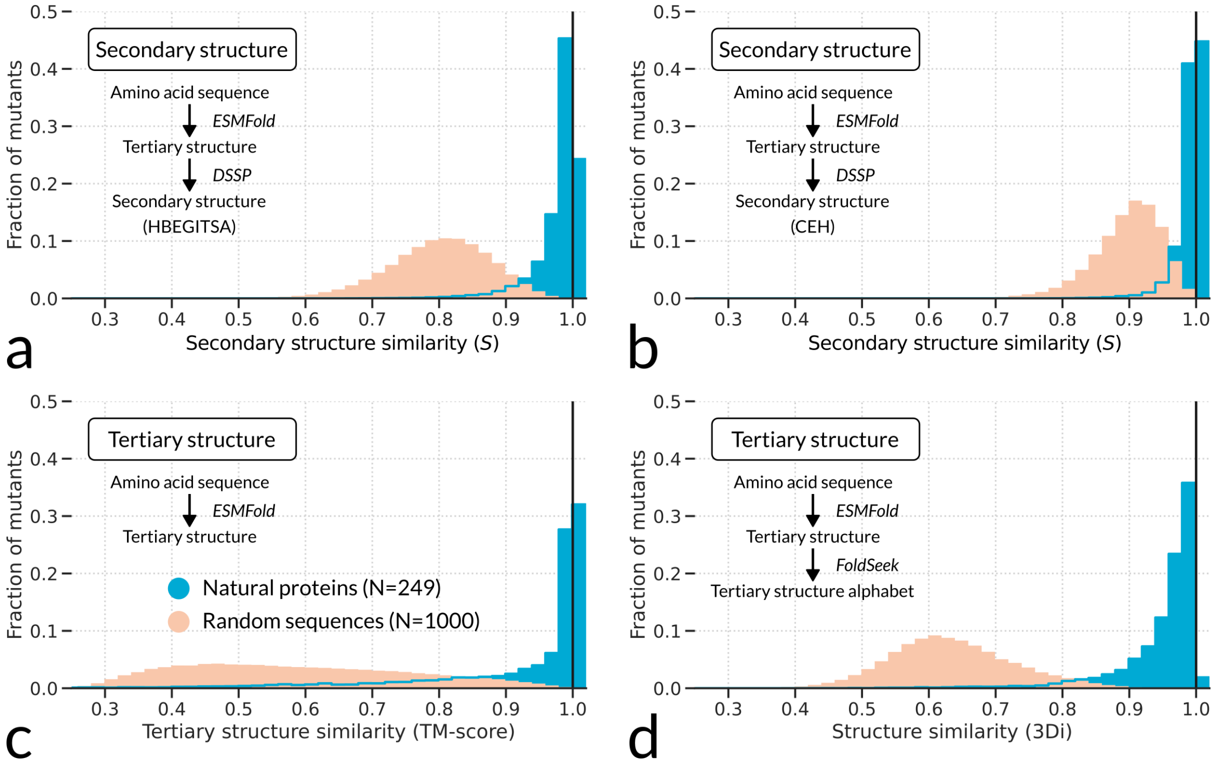


Figure S4 – Mutational profiles of natural and random proteins of length $L = 300$ using different structural representations: (**a**) full DSSP notation (Kabsch and Sander, 1983) distinguishing 8 secondary structure elements (‘`H`’: $\alpha$-helix, ‘`B`’: isolated $\beta$-bridge, ‘`E`’: $\beta$-sheet, ‘`G`’: $3_{10}$-helix, ‘`I`’: $\pi$-helix, ‘`T`’: turn, ‘`S`’: bend, ‘`A`’: disorder), (**b**) standard DSSP notation with three secondary structure elements, (**c**) tertiary structure (alignment of atomic coordinates using TMalign; Zhang and Skolnick, 2005), and (**d**) 3Di representation used by FoldSeek (Van Kempen et al., 2024, such that panels **a**, **b** and **d** are strings). According to most representations except 3Di, natural protein structures feature a high proportion of neutrality $\lambda$. Random sequences always feature low neutrality and are distinct from natural protein structures.

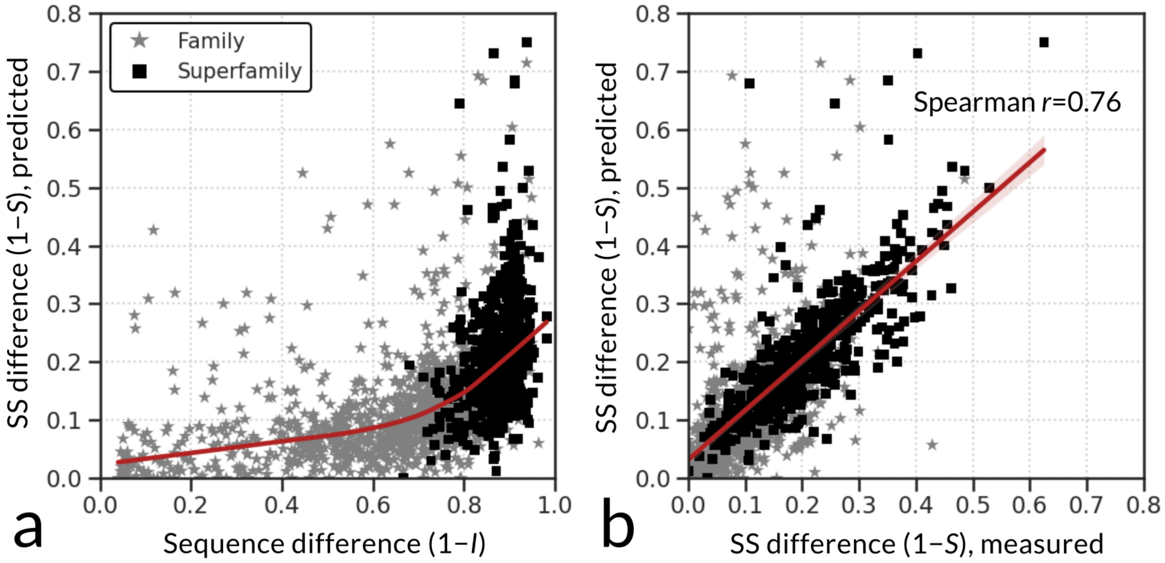


Figure S5 – Correspondence between predicted and measured structural differences across a wide range of sequence differences, *i.e.* for 1594 of 1899 protein pairs from Illergård et al. (2009). (**a**) Predicted differences in secondary structure increase slowly with sequence differences, closely matching the trend in the original study. For each pair, structural alignments from Illergård et al. (2009) were used, in which "non-core" sites which have a gap in one of the sequences are ignored. Data is visualized as in Fig. 2a of Illergård et al. (2009), except that on the x-axis we plot simple sequence difference (fraction of different amino acids) as opposed to inferred substitutions per site. Red line shows the Lowess regression fitted to all data (Cleveland, 1979). (**b**) Structural differences predicted by ESMFold correlate strongly with those measured from the experimentally refined structures. Red line shows linear fit to all data.

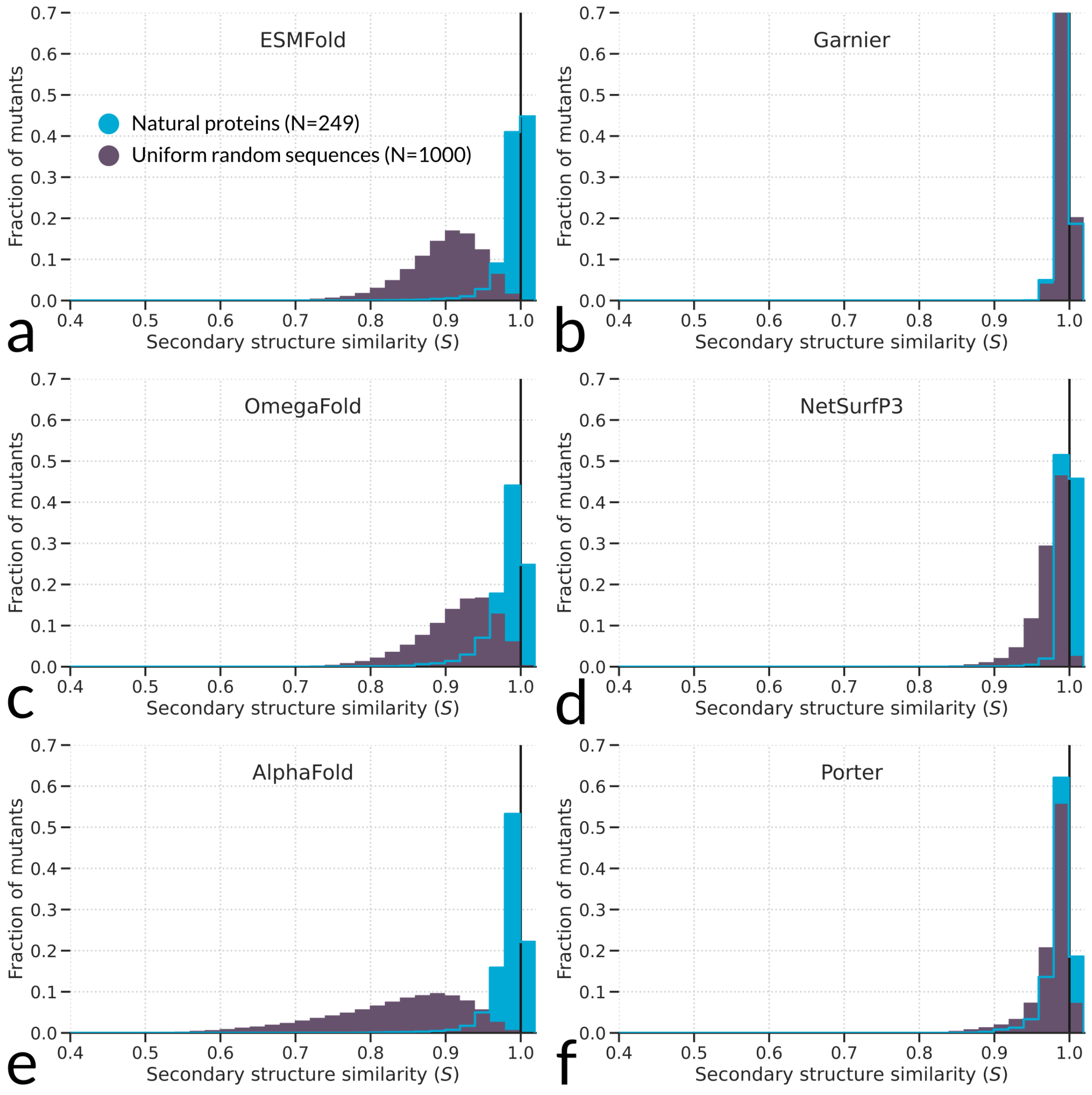


Figure S6 – Mutational profiles of natural and uniform random amino acid sequences (see Fig. S1) of length $L = 300$ predicted by various structure prediction algorithms: (**a**,**c**) ESMFold (Lin et al., 2023) and OmegaFold (Xu et al., 2021), two machine-learning based algorithms designed for one-shot (single-sequence) tertiary structure prediction; (**e**) AlphaFold (Jumper et al., 2021; Kim et al., 2024), a machine-learning based algorithm that relies on multiple sequence alignments to predict tertiary structure; (**b**) Garnier (Garnier et al., 1978), a simple and old algorithm that exploits local sequence contexts to infer the most likely secondary structure type for each residue; (**d**,**f**) NetSurfP3 (Høie et al., 2022) and Porter (v5 Torrisi et al., 2019), two machine-learning based algorithms for secondary structure prediction, with NetSurfP3 designed for one-shot prediction and Porter5 relying on multiple sequence alignments. All algorithms predict narrow and highly robust mutational profiles for natural proteins. Tertiary structure prediction algorithms (**a**,**c**,**e**) consistently predict low robustness for uniform random protein sequences, pointing to a strong evolutionary signature in natural protein structures. By contrast, secondary structure prediction algorithms (**b**,**d**,**f**) predict high robustness for uniform random sequences, which may reflect poor internalization of protein folding (see also Fig. S7).

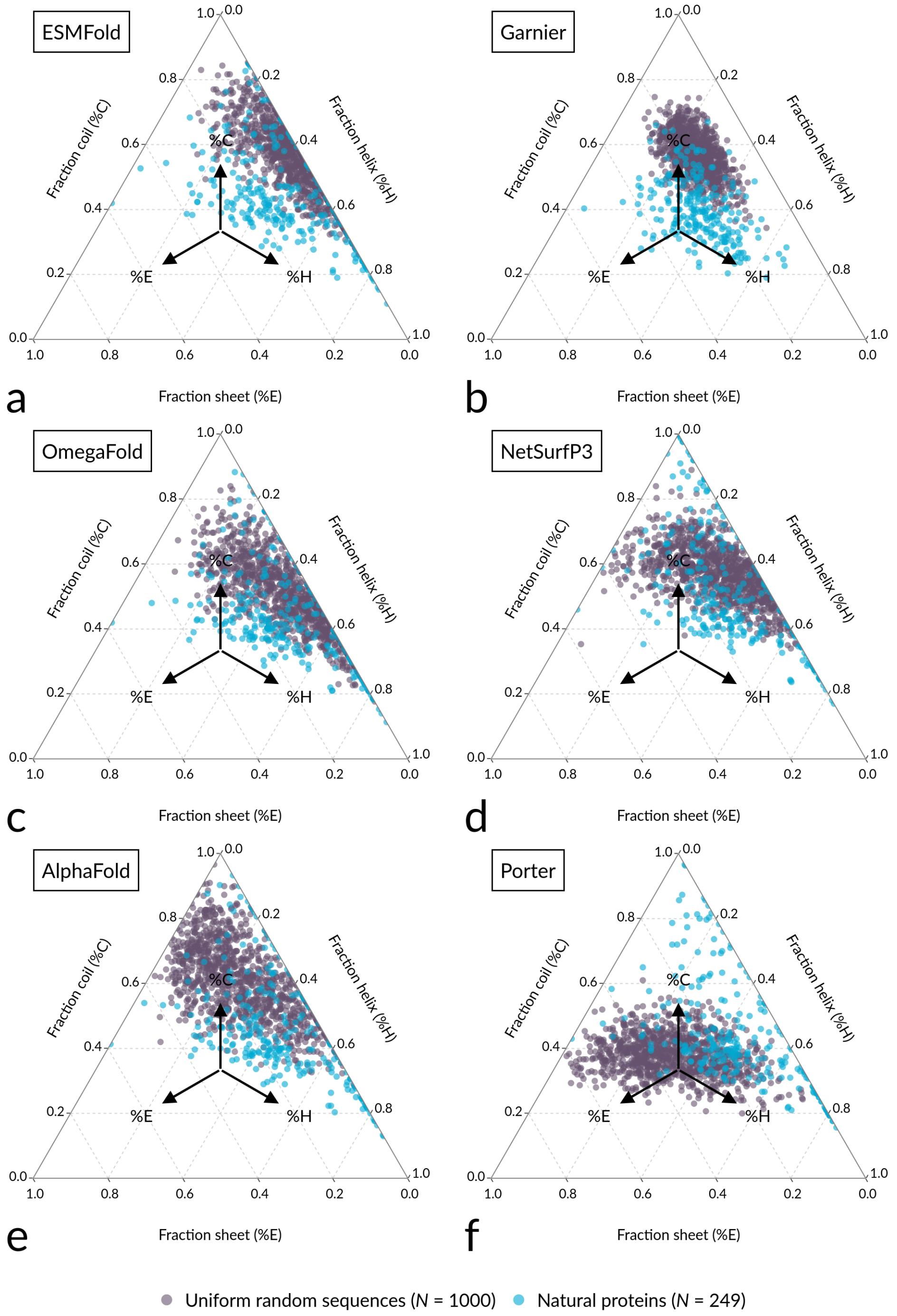


Figure S7 – Fractions of coil, $\alpha$-helix and $\beta$-sheet among secondary structures of natural and uniform random protein sequences, predicted by different algorithms (see Fig. S6). While there is no reference golden standard for secondary structure composition of random, let alone natural proteins, these data provide a glimpse into the inner workings and biases of different algorithms. Most algorithms predict a similar wide distribution of secondary structure compositions for natural proteins. For uniform random sequences, they show larger differences. Predictions by Garnier (**b**) exhibit low variability in secondary structure composition, with not a single protein that completely lacks $\alpha$-helices or $\beta$-sheets. Structure predictions of uniform random sequences by Porter (**f**) exhibit high fractions of $\beta$-sheet, contrasting with previous studies which have suggested that $\alpha$-helices should be more prevalent (Minervini et al., 2009; Tretyachenko et al., 2017; Heames et al., 2023), although one study has claimed the reverse (Yu et al., 2016). AlphaFold (**e**), which like Porter is not designed for structure prediction from single sequence, showcases relatively low levels of $\alpha$-helix. ESMFold (**a**) shows a low prevalence of $\beta$-sheets, whereas OmegaFold and NetSurfP3 (**c**,**d**) show similar composition distributions with slightly higher prevalence of $\beta$-sheets than ESMFold.

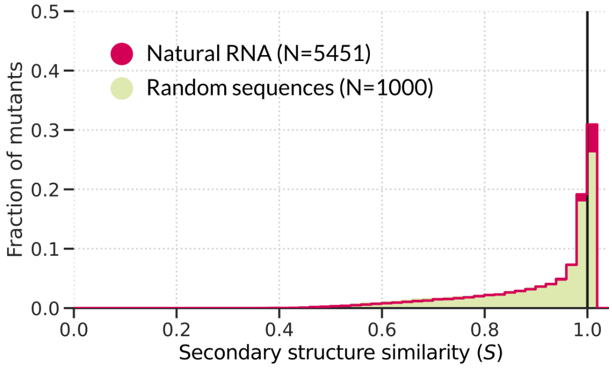


Figure S8 – Machine-learning based RNA secondary structure predictions by MXfold2 (Sato et al., 2021) recover the broad mutational profile of RNA, as well as the strong overlap between natural and random sequences that are described in the main text.

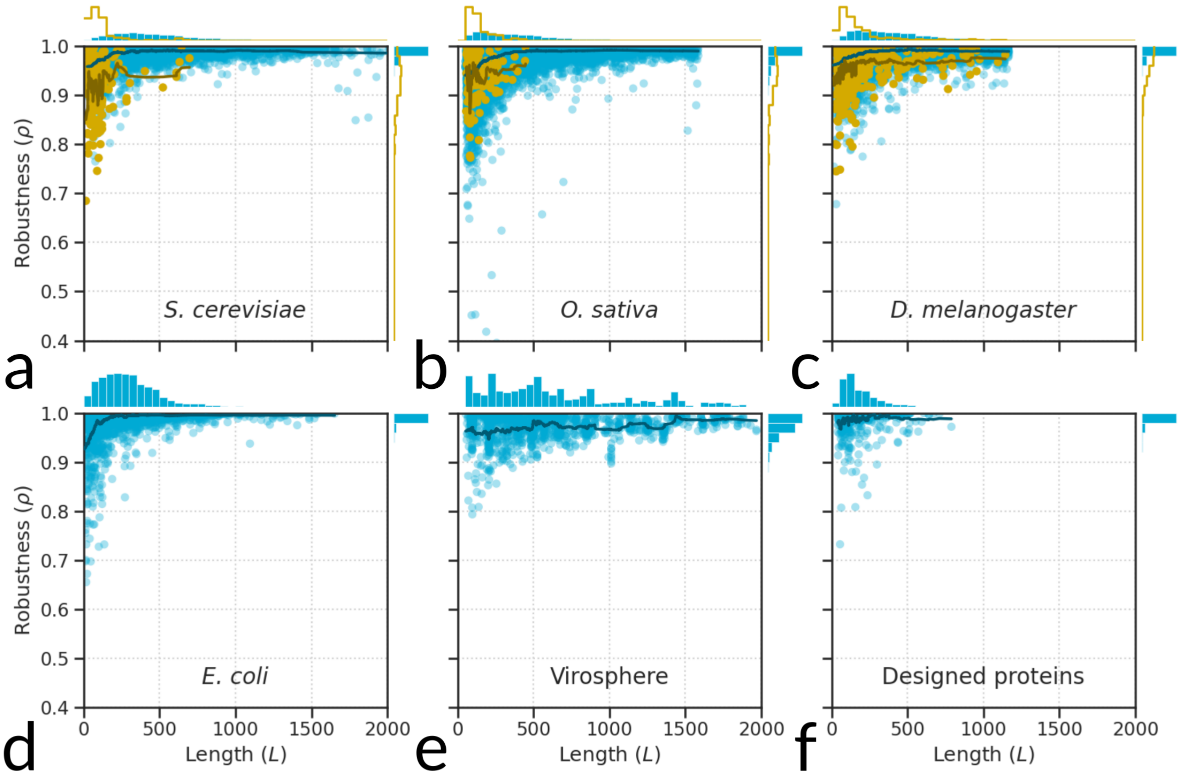


Figure S9 – Predicted mutational robustness versus sequence length across six big datasets, revealing (1) high robustness throughout the tree of life, (2) low robustness for short proteins, and (3) low robustness for *de novo* proteins even when corrected for their short sequence length. (**a**–**c**) Proteomes of three eukaryotic species with *de novo* annotations, *Saccharomyces cerevisiae*, *Oryza sativa* and *Drosophila melanogaster* (Vakirlis et al., 2020; Chen et al., 2024; Peng and Zhao, 2024). For *O. sativa* and *D. melanogaster*, only proteins up to length $L < 1585$ ($N = 19,436$) and $L = 1167$ ($N = 13,094$), respectively, were assessed due to computational limitations. (**d**) *Escherichia coli* proteome ($N = 4295$). (**e**) All viral proteins from UniProt with the maximum annotation score ($N = 1430$; uni, 2025). (**f**) All proteins from the Protein Design Archive (PDA, $N = 952$; Chronowska et al., 2025). As in Fig. 4 in the main text, running means are displayed separately for *de novo* and established proteins.

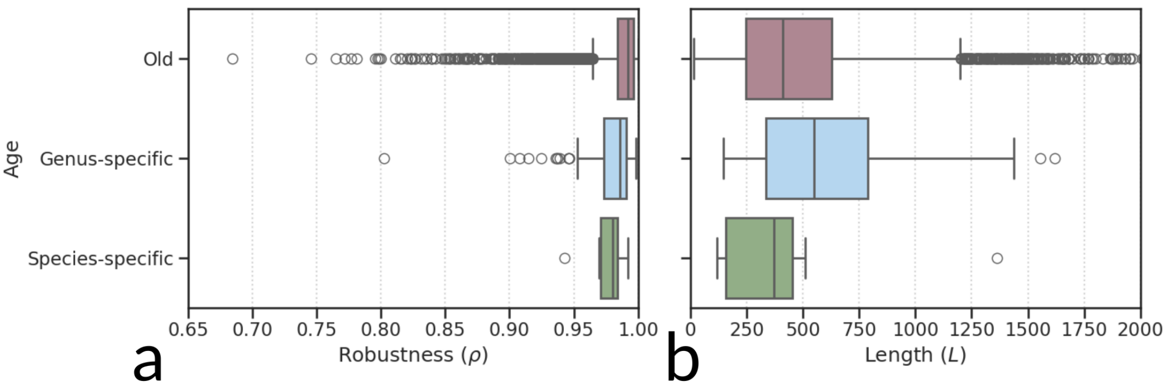


Figure S10 – Phylostratigraphic analysis of *S. cerevisiae* proteome, using annotation from (Ekman and Elofsson, 2010), shows that (**a**) mutational robustness is lowest in young and highest in old proteins. (**b**) Young proteins also tend to be relatively short. Note that the sample sizes are vastly different: we have 7 species-specific, 140 genus-specific, and 5258 "old" proteins (any protein absent from the annotated data of Ekman and Elofsson, 2010).
